# Vertically inherited microbiota and environment modifying behaviors conceal genetic variation in dung beetle life history

**DOI:** 10.1101/2024.01.17.576108

**Authors:** Patrick T. Rohner, Armin P. Moczek

**Affiliations:** Department of Ecology, Behavior, and Evolution, University of California San Diego, La Jolla, CA 92093, United States; Department of Biology, Indiana University Bloomington, IN 47405, United States

**Keywords:** Heritability, evolvability, plasticity, host-microbiome interactions, organism-environment interactions, developmental niche construction

## Abstract

Diverse organisms actively manipulate their (sym)biotic and physical environment in ways that feedback on their own development. However, the degree to which these processes affect microevolution remains poorly understood. The gazelle dung beetle both physically modifies its ontogenetic environment and structures its biotic interactions through vertical symbiont transmission. By experimentally eliminating i) physical environmental modifications, and ii) the vertical inheritance of microbes, we assess how environment modifying behavior and microbiome transmission shape heritable variation and evolutionary potential. We found that depriving larvae from symbionts and environment modifying behaviors increased additive genetic variance and heritability for development time but not body size. This suggests that larvae’s ability to manipulate their environment has the potential to modify heritable variation and to facilitate the accumulation of cryptic genetic variation. This cryptic variation may become released and selectable when organisms encounter environments that alter the degree to which they can be manipulated. Our findings also suggest that intact microbiomes, which are commonly thought to increase genetic variation of their hosts, may instead reduce and conceal heritable variation. More broadly, our findings highlight that the ability of organisms to actively manipulate their environment may affect the potential of populations to evolve when encountering novel, stressful conditions.

## Introduction

Symbiotic microbial communities emerge as a critical factor in the development and evolution of their hosts [1–4]. From a microevolutionary perspective, these interactions are especially significant when symbionts are vertically transmitted from one generation to the next. In these cases, standing genetic variation and responses to selection may not only depend on the host’s genetic makeup but also that of its inherited symbionts as well as the interactions between the two. While there is increasing evidence that microbiomes shape host evolution and development [e.g., 5, 6, 7], the effects of symbionts on phenotypic and genetic variation of their host remains poorly understood [8], especially in cases where symbiont communities are complex and where hosts actively manipulate their environment, thereby influencing presence and function of symbionts. Here, we study the effects of microbiomes and the environment modifying behaviors of their hosts on microevolutionary processes.

Microbial symbionts can affect heritable variation of their hosts in a variety of ways [8]. For instance, the presence of microbes may increase heritable variation if the microbial communities that are vertically transmitted in different host lineages themselves vary in composition and in the phenotypic effects they have on their respective hosts [9, 10]. In these cases, similarity (or dissimilarity) in host phenotype expression may be a function of shared (or divergent) microbial communities. Microbiomes may thus increase genetic variation in host populations and provide added substrate for selection to act upon. However, if symbionts are faithfully inherited over evolutionary timescales, hosts may evolve to become reliant on their microbiomes [e.g., by outsourcing key processes: 11, 12], or conversely, where symbionts are critical to accessing otherwise recalcitrant resources hosts may evolve inheritance mechanisms that increase symbiont fidelity [e.g., vertical transmission and environmental filtering: 13, 14]. In such cases, microbiomes may evolve to become critical components of normative host development and requirements for robust trait expression [15]. If so, the contributions of microbiomes to host development may increase the host’s ability to buffer against deleterious environmental and genetic perturbations [i.e., developmental capacitance: 16]. Likewise, the loss of symbionts may cause environmental stress. Intact host-symbiont relationships may thus also promote the robustness of phenotype expression, potentially shielding (cryptic) genetic variation [17–19] form being exposed to selection. The role of microbiomes in host genetic variation and evolutionary potential may thus be manifold and complex. Here, we use a quantitative genetic approach to assess how the presence (or absence) of microbial communities shapes heritable variation within a host population.

Exactly which microbial taxa engage with a given host may also be in part influenced by host behavior and morphology [20]. Many animals have evolved properties that facilitate the assembly, transmission, and maintenance of their symbiont communities, such as the inheritance of intracellular bacteriocytes (e.g., in aphids [21]), the development of specific organs that facilitate the assembly and function of microbial communities (e.g., the development of the light organ in the Hawaiian bobtail squid [22]), or the construction of external environments that benefit microbial communities. For instance, cockroaches and termites engage in behaviors that, on the colony level, ensure sharing of symbionts and reinoculation across molts [23]. Similarly, *Nicrophorus* carrion beetles use various parental care behaviors to ensure that their offspring are predominantly colonized by maternal (rather than environmental) bacteria [24], and fungus gardening ants maintain microbial taxa in cuticle pockets that prevent the invasion of competing fungi [25]. These examples highlight that hosts can, via their development and behavior, influence which microbes they associate with and thus the nature of interactions with them. However, how such environment modifying behaviors shape the effects microbes have on host heritable variation and evolvability is poorly understood. Here we use dung beetles, their environment modifying behaviors, and their vertically inherited complex microbial communities to jointly investigate the roles played by microbiomes and host behaviors in shaping genetic variation in host life history.

Onthophagine dung beetles are uniquely suited to study the contribution of microbiomes and environment modifying behaviors of their hosts to microevolutionary processes. Females of many species construct underground chambers filled with processed and compacted cow dung [26]. In each of these ‘brood balls’, females deposit a single egg. During oviposition, mothers place each egg onto a small mount of their own excrement, the so-called “pedestal”, representing a microbial inoculate that is consumed by the larva upon hatching. In so doing, the mothers’ gut microbiome is transmitted vertically to its offspring [27]. These vertically transmitted microbial communities have been shown to be host species- and population-specific [28] and to yield deleterious fitness consequences if withheld [7, 29-31]. In addition to the vertical inheritance of gut microbes, larvae also physically modify their brood ball by continuously feeding on its content, excreting back into their brood ball, spreading excreta, and re-eating the increasingly modified composite [7, 27]. As the developing larva continually defecates, works its own excrement into the brood ball, and then re-eats the resulting mixture, the maternally inherited gut microbiome is spread throughout the brood ball, thereby increasing its ability to pre-digest macromolecules outside the larval gut, at least as assayed by *in-vitro* studies [7, 32]. Experimental withholding of these modifications results in prolonged development, smaller size, and reduced secondary sexual trait expression [but see: 33], suggesting that these environmental modification aids in the extraction of nutrients from an otherwise recalcitrant diet and thus feeds back onto larval development [32, 34]. The brood ball can thus be regarded as an extended phenotype [35], or as a product of maternal and larval niche construction [36]. However, whether and how the interactions between maternal microbiota and host behaviour impact standing genetic variation residing within host populations remains unknown.

Here, we assess the role of vertically inherited microbiomes and their interactions with environment modifying host behaviors in shaping genetic variation in the dung beetle *Digitonthophagus gazella*. Combining a quantitative genetic breeding design with an experimental elimination of i) physical modifications of the environment and ii) the vertical inheritance of microbial symbionts, we assess how microbial communities and their cultivation by their host shape heritable variation. Our findings suggest that the presence of ontogenetic environmental modifications and vertically inherited symbionts may conceal otherwise cryptic genetic variation and thus impact heritable variation visible to selection. Taken together, our findings emphasize the potential of the interactions between hosts, their microbes, and the environment to shape microevolutionary dynamics.

## Methods

### General laboratory rearing and experimental manipulations

*Digitonthophagus gazella* (Fabricius, 1787) was collected in March 2021 near Pretoria, South Africa, sent to Indiana University, Bloomington, USA and kept under standard laboratory conditions [e.g., 37, 38]. To obtain laboratory-reared F1 individuals, we repeatedly transferred 4 to 6 wild-caught (F0) females from the laboratory colony into rectangular oviposition containers (27cm × 17cm × 28cm) filled with a sterilized sand-soil mixture and topped off with ca. 800g defrosted cow dung. After 5 days, brood balls were sifted from the soil and kept in plastic containers filled with soil at constant 29°C.

Newly emerged F1 offspring were kept in single-sex containers for at least 7 days at 26°C. Thereafter, 30 males (sires) were housed together with 3 females (dams) each in separate containers equipped with sterilized soil and defrosted cow dung for at least 4 days (see fig. S1). Females were then individually transferred to oviposition containers (27cm × 8cm × 8cm) filled with a sterilized sand-soil mixture and topped off with 200g defrosted cow dung [see 39] and kept at 29°C. Brood balls were collected after 5 days. We reared the F2 offspring in standardized, artificial brood balls as described previously [40]. In brief, we opened all natural brood balls and transferred eggs individually into separate wells of a standard 12-well tissue culture plate provisioned with 2.9 (±0.1) grams of previously frozen cow dung. To minimize variation in dung quality and quantity among wells, we thoroughly homogenized a large quantity of cow dung using a hand-held electric cement mixer (Nordstrand, PWT-PM0) prior to the start of the experiment. We only used dung from hay-fed cows, which is less nutritious compared to dung from grass-fed cows [41]. Plates were kept at 29°C and checked for hatching every 24 hours. All F2 were subjected to two fully factorial manipulations of a larva’s ability to shape its (sym)biotic and physical ontogenetic environment:

#### Microbiome manipulation

To manipulate the vertical transmission of microbial symbionts, half of all eggs were surface-sterilized with 200μl of a 1% bleach and 0.1% Triton-X 100 solution, followed by two rinses with deionized water [see 7, 29, 42]. Eggs in the control treatment were rinsed with deionized water only. Eggs were then placed in an artificial, standardized brood ball, either with (‘*intact microbiome transmission’*) or without (‘*disrupted microbiome transmission‘*) the extracted maternal pedestal. The latter was removed from the natural brood ball and transferred into the artificial brood ball using a flame-sterilized spatula [as in e.g., 31]. Note that the bleaching treatment is a standard approach in dung beetles [7, 31] and other taxa (water fleas: [43], tephritid fruit flies: [44]) and there is no evidence for any deleterious effect on embryonic or postembryonic beetle development in dung beetles [7, 31]. Note, however, that although bleach only sterilizes the egg surface and does not come into contact with the beetle embryo, we cannot completely rule out that bleach treatments may have any previously undetected minor effects on postembryonic development besides the disruption of host-symbiont interactions.

#### Manipulation of larval environment modifying behaviour

The capacity of larvae to manipulate their brood ball was experimentally hampered by relocating individuals into a new artificial brood ball 4, 7, 10, and 13 days after eggs were initially transferred using featherweight forceps [see: 32, 34]. This procedure exposes the developing larva repeatedly to new, unprocessed cow dung and prevents the accumulation of environmental modifications applied to the brood ball (‘*impaired brood ball modification’*). Specifically, this procedure prevents larvae from repeatedly feeding on and restructuring dung particles within their brood ball. Relocation into a new brood ball also disrupts their association with the established microbial communities in the previous brood ball. In the control treatment, larvae were allowed to complete their development in their original well. To account for the potential stress induced by repeatedly relocating larvae into new wells, larvae were removed from their brood ball, held with featherweight forceps for approximately 3 seconds, and placed back in their original well 4, 7, 10, and 13 days after eggs were transferred into new plate (‘*intact brood ball modification’*).

Individuals were checked daily to assess juvenile survival and egg-to-adult development time. Individuals were classified as adult on the day they emerged from the pupal cuticle. We also imaged the adult thorax using a Scion camera mounted on a Leica MZ 16 stereomicroscope and measured pronotum width (a suitable estimate for body size, see [45]) using tpsDig2 [46].

### Statistical analysis

To assess the fixed effects of both experimental manipulations on egg hatching success and juvenile survival, we used generalized linear mixed models with binomial error structure in lme4 [47]. Dam nested within sire as well as the 12-well plate individuals were reared in were added as random effects. We used linear mixed models with the same design to test for sex-specific effects on logarithmized pronotum width and development time. We added sex and all interactions with fixed effects to the model. Because sex can only be determined in late larval development [37] sex could not be included in the models for juvenile survival and hatching success.

To test whether our manipulations of the microbiome and larval brood ball modification affected genetic variation and heritabilities individually or in combination, we computed treatment-specific variance components using “animal models” in ASReml-R [48, 49]. ‘Animal models’ are a type of mixed-effects models that have been widely applied to estimate quantitative genetic parameters because they are based on pedigrees rather than strict breeding designs (e.g., [50–52]). In essence, instead of relying on the variance among genetic groupings (e.g., sires), animal models fit the genetic variance component directly based on a relationship matrix (i.e., a matrix summarizing the pairwise relatedness among all individuals) in a linear model with reduced maximum likelihood (REML) (see [49, 53]). Animal models better accommodate unbalanced data and can use information on multiple generations for the estimation of genetic parameters. We used animal models rather than sire models mainly because they allow to estimate additive variances directly [53]. We estimated separate additive and residual variance components for all treatment combinations simultaneously (i.e., intact microbiome transmission and brood ball modification (control); disrupted microbiome transmission; disrupted brood ball modification; or disrupted microbiome transmission and brood ball modification). Sex, treatment, and their interaction were added as fixed effects. The 12-well plates individuals were reared in were added as a random effect. To test whether partitioning the additive and residual variances among treatments significantly increase model fit, we used Likelihood Ratio Tests (LRTs) to compare the full model to one that did not include treatment-specific additive or residual variances. When the overall model indicated changes in variances across treatment combinations, we also conducted pairwise comparisons between treatment combinations. Variances were left unconstrained in all models. Narrow-sense heritabilities (*h^2^*) were computed by dividing the additive genetic variance by the total phenotypic variance in the respective treatment. Evolvability (*I_A_*, i.e., mean-scaled additive genetic variances [54]) were calculated by dividing the treatment-specific additive genetic variances by the square of the treatment-specific mean trait values.

## Results

### Effect of microbiome transmission and brood ball modifications on phenotypic variation

Our full-sib/half-sib breeding design resulted in 1,228 eggs produced by 67 females mated to a total of 25 sires. In total, 932 individuals survived to adulthood. Hatching success was higher when eggs were surface sterilized (χ^2^ = 8.85, P = 0.003). Larval survival was higher when larvae were able to manipulate their brood ball (χ^2^ = 96.60, P <.001) but did not depend on the transmission of maternal microbiomes (χ^2^ = 0.40, P = 0.526; table 1). Larvae that were able to physically modify their brood ball also developed faster (χ^2^ = 206.79, P <.001) and grew to larger adult size (χ^2^ = 240.36, P <.001, table S1). The effect on body size was stronger in males, leading to a decrease of sexual size dimorphism when larvae were prevented from manipulating their environment (sex-by-treatment interaction: χ^2^ = 21.37, P <.001, table S1; fig. 2). Withholding the vertically transmitted microbiome also reduced adult body size (χ^2^ = 55.41, P <.001) and delayed adult emergence (χ^2^ = 42.93, P <.001, table S1). This effect was especially strong in females deprived of maternal microbiota that were unable to modify their brood ball (three-way interaction between sex, microbiome transmission, and brood ball manipulation: χ^2^ = 9.09, P = 0.003, table S1). Microbiome transmission and brood ball modifications thus not only shape phenotype expression but do so in an interdependent and sex-specific manner.

**Figure 1:**
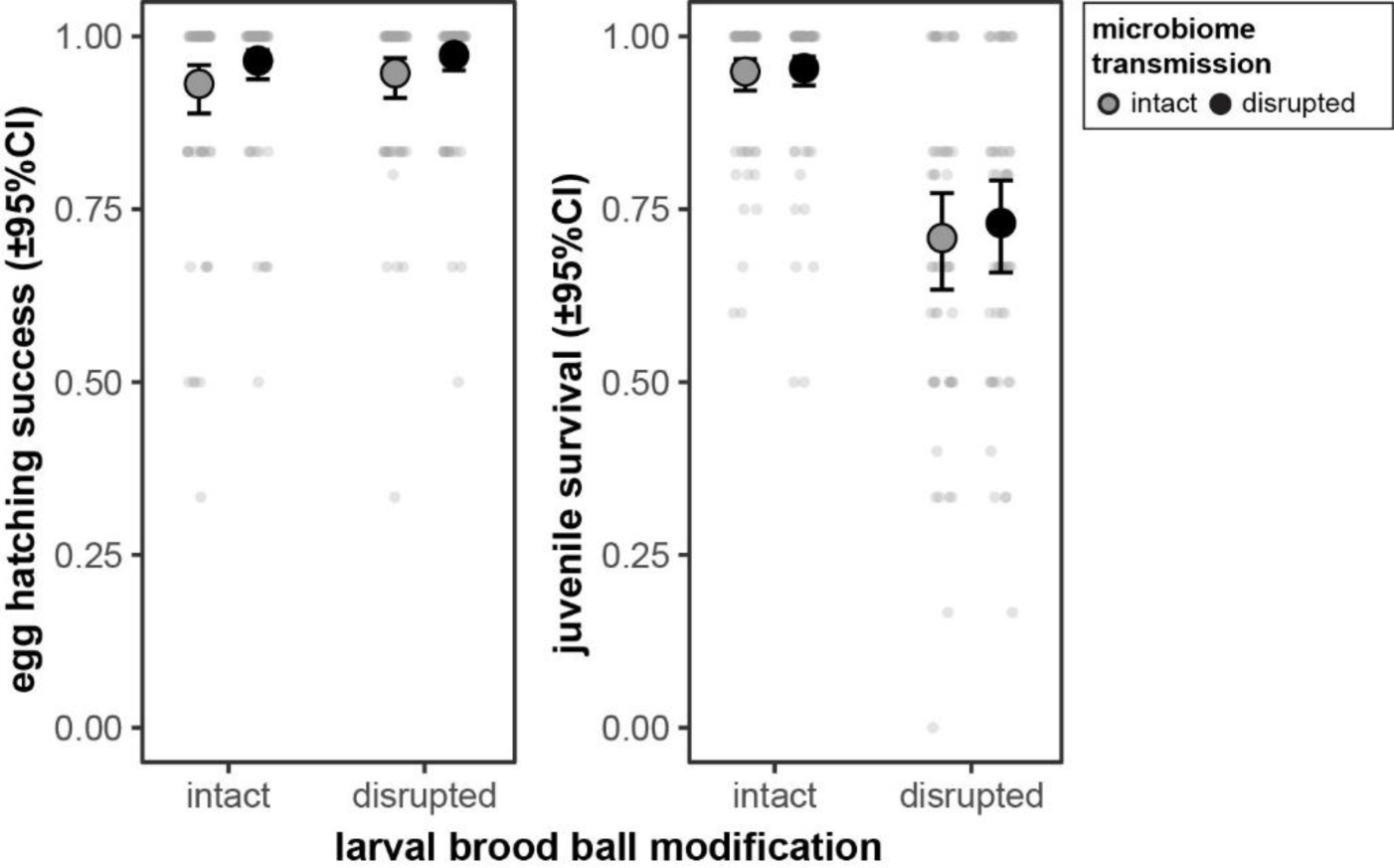
Estimated marginal means and corresponding 95% confidence limits for egg hatching success and juvenile survival as a function of the presence of maternal microbiota and larval brood ball modifications. Individual data points represent treatment-specific full-sib family means.

**Figure 2:**
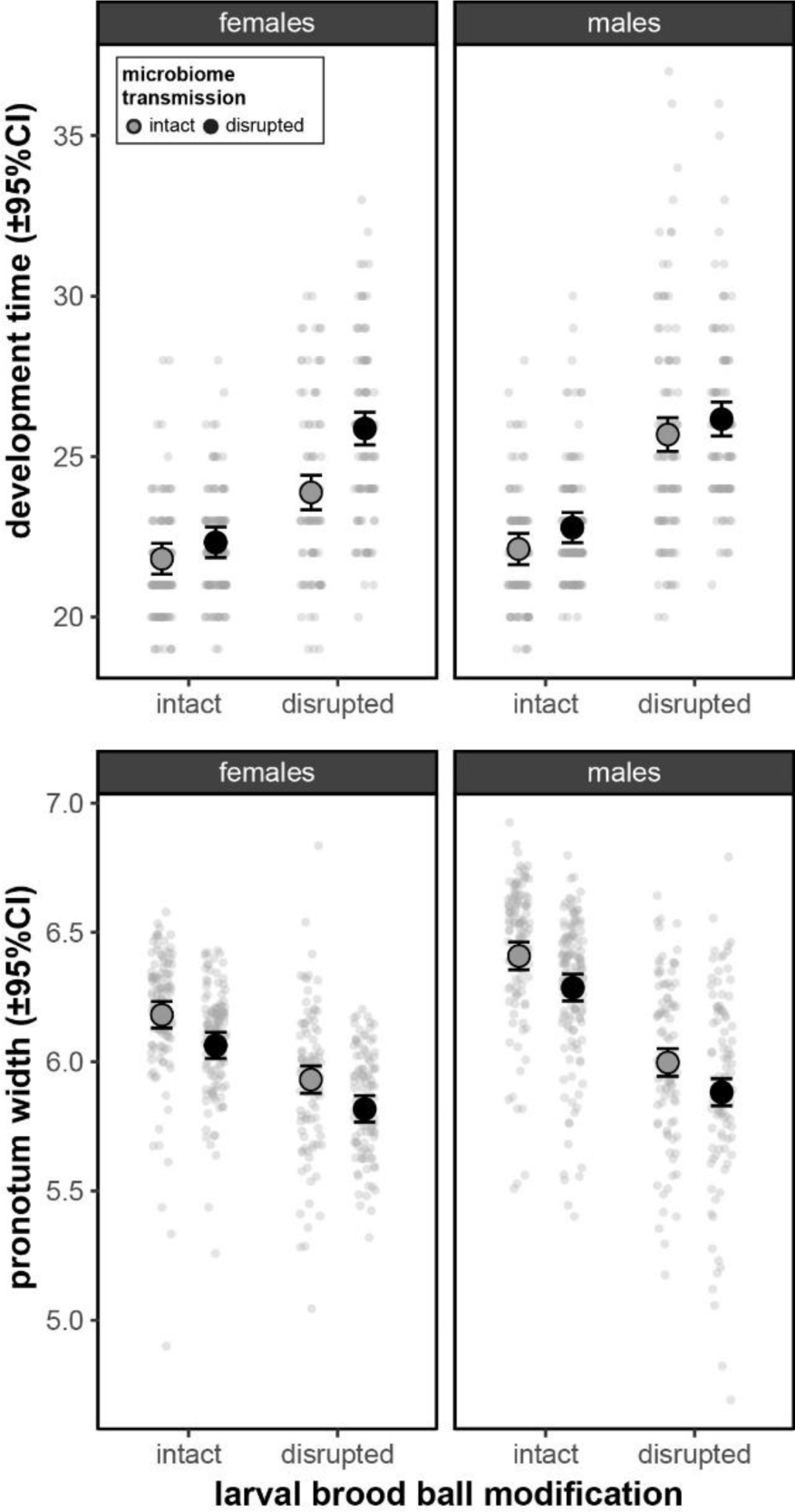
Estimated marginal means and corresponding 95% confidence limits for development time and body size as a function of the presence of maternal microbiota and larval brood ball modifications. Data points represent individual measurements (total n = 932 individuals).

**Table 1:**
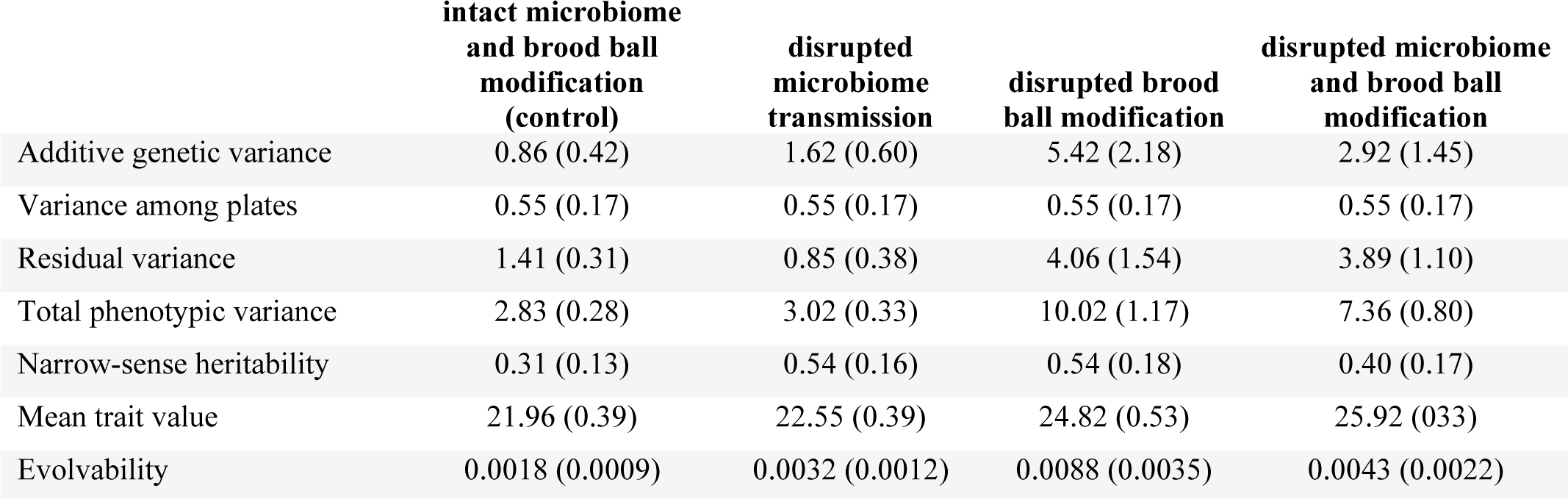
Variance components, heritabilities, trait means, and evolvability (±SE) for development time for the four treatment combinations. Variances were computed using an animal model simultaneously estimating additive and residual variances for each treatment combination. Heritability was calculated by dividing the treatment-specific additive genetic variance by the treatment-specific total phenotypic variance. Evolvability was computed by dividing additive genetic variance by the square of the trait mean.

### Effect of microbiome transmission and brood ball modifications on variance components and heritability

Models including separate additive and residual variances in development time for each of the four treatment combinations fitted the data better than models with no treatment-specific variances (LRT: χ^2^ = 170.9, P <.001, fig. 3, 4), or models that included treatment-specific residual variances only (LRT: χ^2^ = 26.24, P <.001). Pairwise comparisons of variance components across treatments further revealed that preventing larvae from physically modifying their brood ball greatly increased the additive as well as the residual variance in development time and led to an increase in narrow-sense heritability from 0.31 ±0.13 to 0.54 ±0.18 (fig. 3, 4, table 1). Preventing larvae from receiving a microbial inoculate caused a modest increase in the additive genetic variance but decreased the residual variance, leading to an increase in heritability to 0.54 ±0.16. Simultaneously removing brood ball modifications as well as microbiome transmission also led to an increase in the additive and residual variances, and an increase in heritability (*h^2^*= 0.40 ±0.17). Evolvability in the control treatment was low (*I_A_* = 0.0018) but increased considerably when limiting larvae’s ability to shape their biotic (*I_A_* = 0.0032), physical (*I_A_* = 0.0088), or both components of the environment simultaneously (*I_A_* = 0.0043; see table 1).

**Figure 3:**
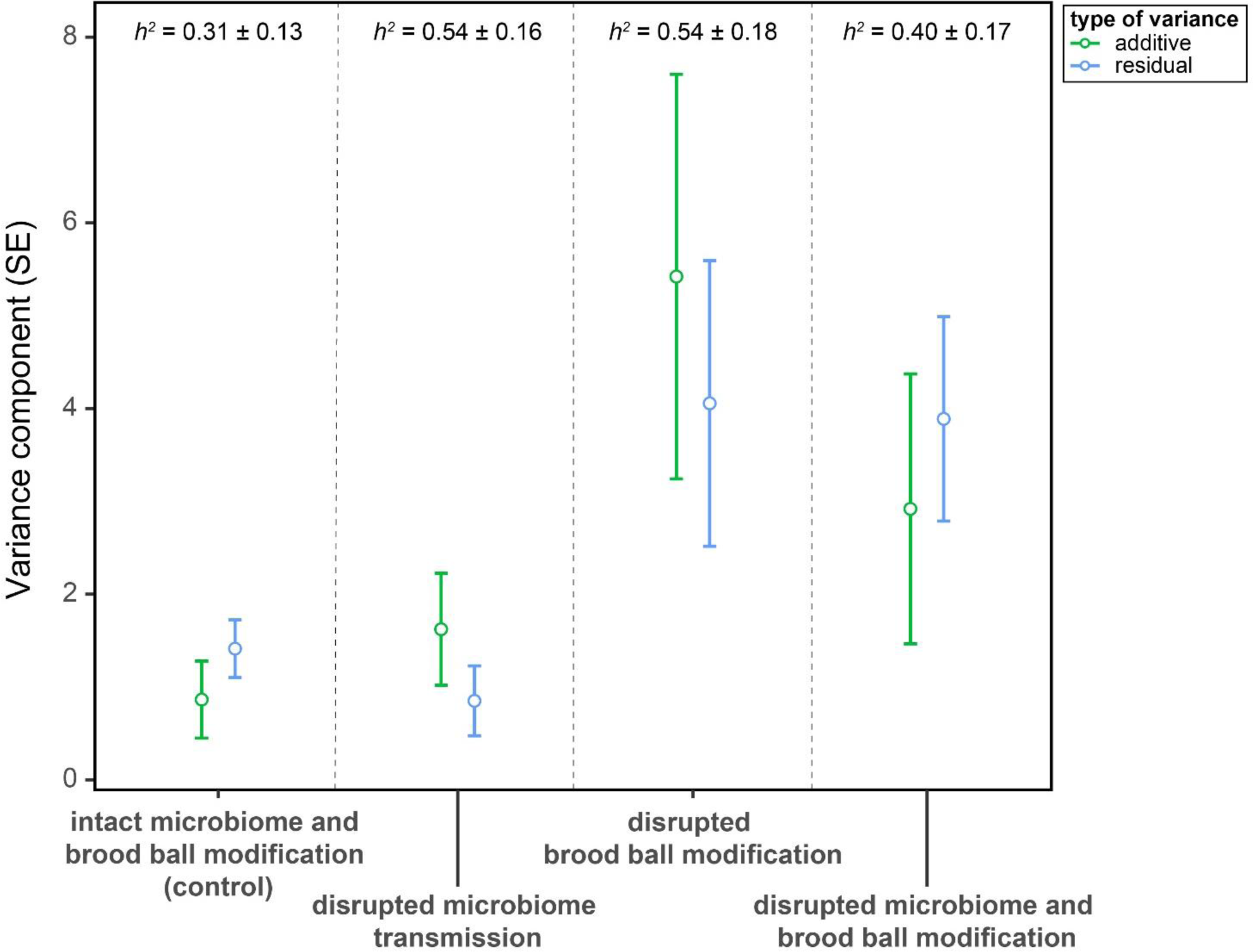
Additive and residual variance components for development time for each treatment combination derived from animal models (total n = 932 individuals). Heritabilities were calculated by dividing the additive variance by the total phenotypic variance (including the variance attributable to the 12-well plate individuals were reared in, see table 1).

Body size also showed significant levels of genetic variation (*h^2^* = 0.64 ±0.11, *I_A_ =* 0.0016 ±0.0004, P <.001). However, in contrast to development time, there was no evidence for treatment-specific additive or residual variances (all P >.900). Using animal models with binomial error structure, we did not find significant levels of additive genetic variation for juvenile survival and egg hatching success (all P >.900).

## Discussion

Using an experimental manipulation of microbiome transmission and dung beetle larvae’s ability to physically modify their brood ball, we empirically assessed the role of symbionts and environment modifying behaviors in shaping phenotypic and heritable variation. Experimentally eliminating microbiome transmission and brood ball modification generally led to an increase in additive genetic variance in development time (figs. 3 and 4). This caused an increase in the evolutionary potential as quantified by heritability (the proportion of the total variance that is additive) and evolvability (expected proportional change under a unit strength of selection [54]). This is consistent with the hypothesis that host-symbiont associations and environment modifying behaviors reduce environmental stress and promote developmental stability and the accumulation of cryptic genetic variation. Because development time is a major life history trait often involved in local adaptation [e.g., in the related dung beetle *Onthophagus taurus*, 39], brood ball modifications and host-symbiont relationships may thus have the potential to influence a population’s ability to respond to selection, especially when encountering novel environments. However, these effects were only found for development time while heritable variation in body size was independent of a larva’s ability to manipulate its environment. Taken together, our data suggest that the interactions between developing larvae and their ontogenetic environments have the potential to contribute to microevolutionary dynamics in one of two traits found to exhibit heritable variation.

**Figure 4:**
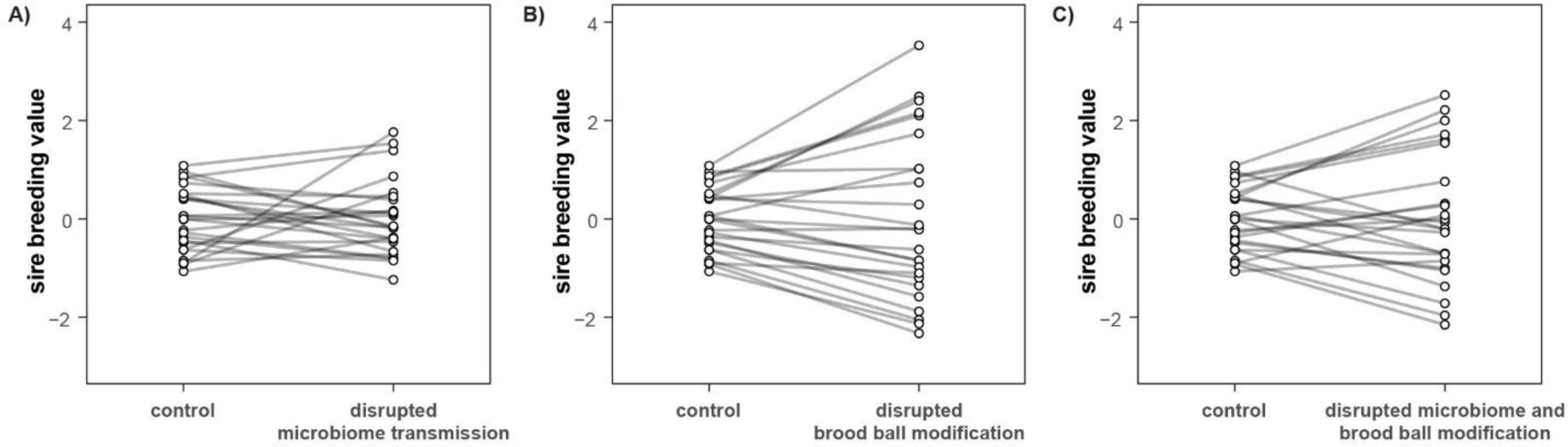
Genetic breeding values (genetic merit) for development time of 25 sires across control and manipulated environments. Lines indicate the nge in genetic values of each sire when microbiomes and/or brood ball modifications are disrupted. The large variation in the slope of these reaction ms indicates that genotypes differ in their response to the experimental manipulation. Breeding values represent best linear unbiased predictions LUPs) and were extracted from an animal model including all individuals (n = 932 offspring).

### Microbiome transmission and brood ball modifications reduce phenotypic and additive genetic variation

Many organisms actively modify the (sym)biotic and physical environment they experience [36, 55]. Because environments serve as major developmental regulators [56, 57], modifications made to the ontogenetic environment can feed back onto an individual’s own development and shape developmental outputs. Environmental modifications thus indirectly shape genotype-phenotype maps, especially of those phenotypes that show plastic responses to the modified environmental variable [1]. Because additive genetic variance is environment-dependent [e.g., 58, 59], the presence of symbionts or environment modifying behaviors can also affect heritability and evolvability and, therefore, a population’s potential to respond to selection [60, 61]. By depriving larvae from their maternal microbiome and their ability to manipulate their brood ball, we found an increase in phenotypic variance. These effects were stronger when a larva’s ability to physically manipulate its environment was impeded compared to the removal of maternally transmitted symbionts. These findings contrast with other studies where microbiomes increased host trait variation [8], but are consistent with the idea that the ability to structure their environment increases organisms’ robustness against environmental perturbations. Intriguingly, we also found a disproportionate increase in the amount of additive genetic variance relative to the total treatment-specific phenotypic variance (i.e., *h^2^*) in all three treatment combinations in which larvae were either deprived of their symbionts and/or had their ontogenetic environmental modifications disrupted. This is consistent with the hypothesis that symbiont inheritance and host behaviors, when intact, not only buffer against environmental but also genetic perturbations, thereby enabling the accumulation of cryptic genetic variation. When organisms’ capacity to compensate for stressful environmental conditions becomes limited, this previously cryptic variation can be exposed and become visible to selection [18]. This suggests that, if disturbed, environment modifying behaviors and host-symbiont interactions may act as evolutionary capacitors through the release of previously cryptic genetic variation [15].

Although we eliminated microbiomes and larval behavior experimentally, natural conditions may also limit or compromise larvae’s ability to modify their physical and microbial environment to their advantage. For example, natural and human-mediated range expansions of both dung beetles and dung producers are common [26]. During colonization, adult dung beetles may thus encounter and utilize novel dung types less accessible to their resident microbiome, as for instance in *Onthophagus australis*, a dung beetle native to Australia which switched from marsupial to cow dung upon introduction of cattle to the continent [62]. Even stronger effects may be expected for the large number of dung beetle species that are primarily associated with life stock. For instance, the widespread treatment of cows with antibiotics not only changes the microbial composition of cow dung but also disturbs the microbiome of beetles that feed on contaminated dung [63]. These agricultural practices may hence reduce dung beetles’ abilities to shape their biotic environment and, in the process, release previously accumulated cryptic genetic variation. Similarly, agricultural management practices that change soil or dung composition may impact the extent to which larvae are able to physically manipulate their ontogenetic environment: for instance, compared to grass-fed cows, hay-fed cows produce dung that contains a much greater fraction of coarse fibers. Hay dung resides longer within the larval gut, larvae feeding on it require more time to complete development, and emerge as smaller adults [41]. Nutritional and physical differences between dung types may thus influence the effectiveness of larval brood ball modification behavior.

Taken together, we found that symbionts and environment modifying behaviors may shield genetic variation from manifesting on the phenotypic level and thus remain cryptic. However, while we found an effect of our manipulations on the heritability of development time, we did not find similar effects on body size, suggesting that effects on genetic variation are trait specific. We also found no effect on variance in juvenile survival and hatching success, yet, we did not find evidence for heritable variation in these two traits to begin with which may be due to limited power when estimating variance components in binary response variables. Further research will be necessary to test whether the trait differences between development time and adult size are driven by selection for genetic or developmental integration between environment modifying traits and recipient traits [64, 65]. Similarly, the effect of our manipulations on the microbial community inside the larval gut and the brood ball requires further scrutiny. For instance, the increase of residual variation when microbiomes are withheld could be caused by random colonization of larvae with environmental microbes. Future research using sequencing of microbiota and their function will be required to better understand how the removal of maternal microbial inocula shape offspring microbiomes.

### Genetic variation for the dependence on symbionts and environment modifying behaviors

Previous work indicated differences among species or populations in the effects of brood ball modification and the vertical transmission of microbiomes on dung beetle performance [7, 30, 32]. Here, find that genetic variances differ between treatments, implying that there is genetic variance for responses to the elimination of microbiomes and brood ball modifications (fig. 4; genetic cross-environmental correlations shown in fig. S2). The phenotypic similarity between relatives in a population could thus be explained, in part, by heritable variation for how developing organisms interact with their microbiome or how larvae manipulate their ontogenetic environment. Host-symbiont interactions and environment modifying behaviors may thus indirectly respond to selection and evolve.

While we found heritable variation in the response to the withholding of maternal microbiomes and brood ball modifications, the causes of this variation remain unclear. Genotypes may, for instance, vary in the effectiveness of their behaviors to physically manipulate ontogenetic environments. Yet, there may also be heritable differences in the susceptibility of developing larvae to the environmental conditions generated by the absence of these modifications. Similarly, it is unclear why genotypes differ in their response to the removal of vertically transmitted microbes. Adult mothers differ at least in part in the taxonomic composition of their microbiome as do their offspring [27], raising the possibility that (epistatic) interactions between beetle hosts and the presence of microbial symbionts may shape heritable variation within host populations. However, microbial communities are complex and their patterns of vertical transmission and effect on host fitness are still poorly understood. While the presence of microbiomes clearly enhances host development [7], recent findings suggest that not all vertically-transmitted microbial members are necessarily beneficial [30]. Our finding that eggs with intact microbiomes had a lower hatching success compared to sterilized eggs may also indicate the presence of harmful microbiota that affect early host development, suggesting that microbiome-mediated effects on host fitness are complex. The precise mechanism mediating heritable differences in the response to the removal of environmental manipulations thus remains elusive and requires further investigation.

### Sex-specific interactions between different components of environmental modifications

In addition to treatment effects on genetic variation, we also found previously undocumented interactions between brood ball modifications, the presence of vertically transmitted microbiota, and the sex of the developing beetle larva. These interactions were especially pronounced in the prolongation of development time in females that could neither benefit from vertically-transmitted microbiota or brood ball modifications. Similarly, preventing larvae from conditioning their brood ball reduced sex differences in adult size. Genetic or environmental changes in the interactions between larvae and their ontogenetic environment may thus affect sexual dimorphism, a major aspect of phenotypic variation in this species [45]. Such non-additive effects of microbiome and larval environment modifying behavior are consistent with the hypothesis that the two interact. For instance, while the microbial community inside a brood ball is likely shaped by the presence of vertically transmitted microbiomes, the extent to which this same microbial community is then able to colonize and modify the brood ball may in turn be determined by the activities and physical modifications made by a larva [32, 34]. Similarly, as larval developmental trajectories diverge as a function of sex (e.g., due to costly ovarian differentiation in female but not male larvae; [66]), changes in environmental conditions experienced by larvae may fuel sex-specific responses to experimental or natural alterations of environmental conditions. Given that males and females of diverse insects commonly differ in growth responses to environmental conditions [67, 68], sex-differences in the response to the presence (or absence) of microbiomes or modified ontogenetic environments, as documented here, may thus be similarly widespread.

### Conclusions

Using an experimental reduction of two distinct routes through which developing dung beetles shape their ontogenetic environment, we demonstrate the potential of host-symbiont interactions and environment modifying behaviors in shaping additive variation, heritability, and evolvability for some life history traits but not others. Furthermore, we found heritable variation for the response to the elimination of environmental modifications. Although the mechanisms underpinning these patterns remain elusive, our findings underscore the potential of environment modifying behaviors in shaping heritable variation in populations. This suggests that rather than merely reacting to environmental conditions, organisms may evolve to shape their immediate environments in ways that in turn may feed back to impact their own microevolutionary trajectories. Taken together, these data call for further investigation into the mechanisms by which developing organisms shape their ontogenetic environment, and the conditions under which these interactions may shape microevolutionary outcomes [56, 61, 69].

## Acknowledgements

We thank Christian Descholdt for collecting beetles in South Africa, Anna Macagno for support with beetle husbandry, and Erik Postma, Michael Wade, and Wolf Blanckenhorn for advice on the statistical analysis. We would also like to thank the Associate Editor and two anonymous Reviewers for their constructive and helpful comments.

## Author Contributions

P.T.R. and A.P.M. conceived and designed the study; P.T.R. collected all data and performed all analyses; P.T.R. and A.P.M. drafted the initial version of the manuscript and contributed to later versions of the manuscript.

## Data Availability

All data underlying this study will be made publicly available on Dryad.

## Supplementary material

**Figure S1:**
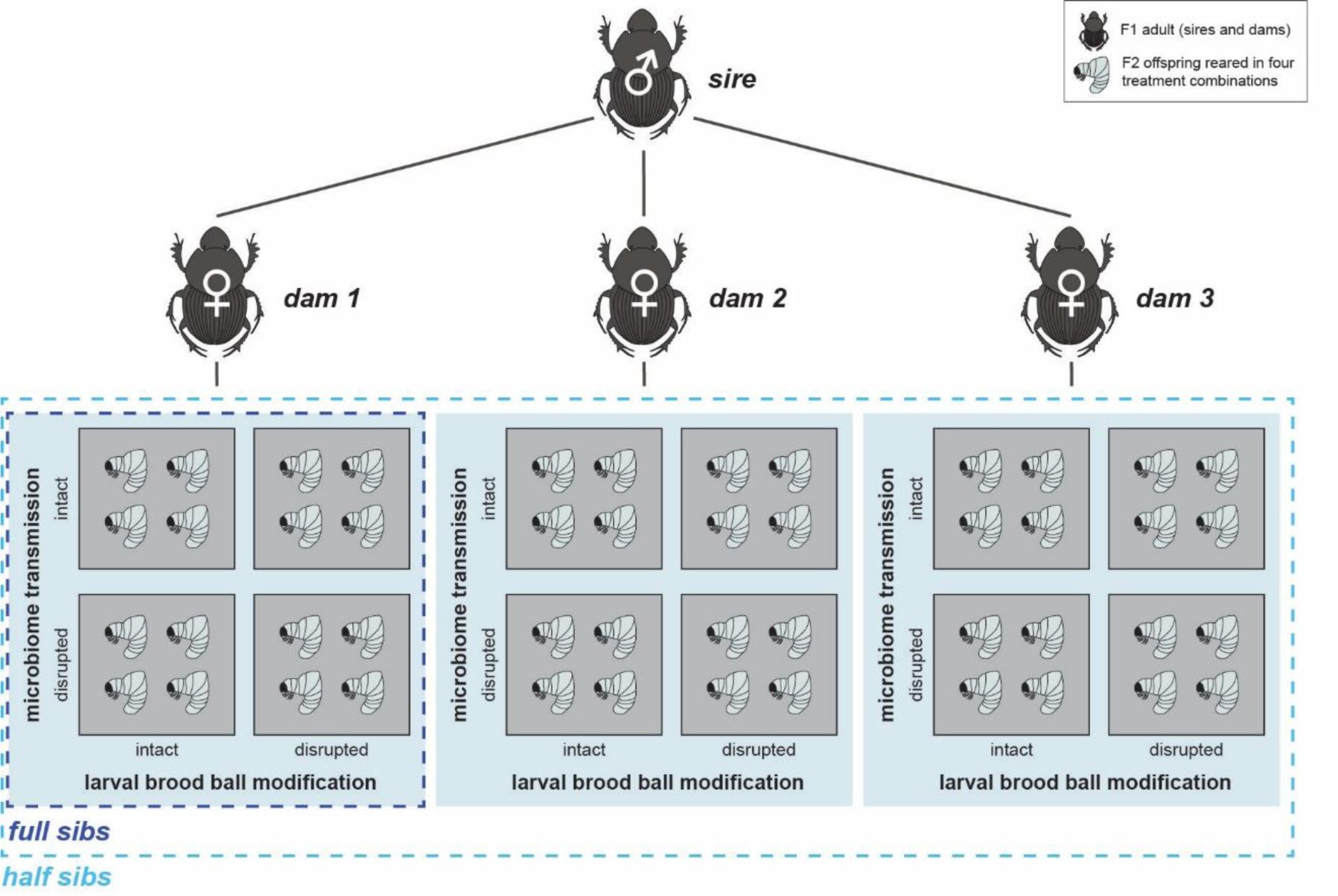
To generate offspring with varying levels of relatedness for the estimation of additive genetic variances, we reared 30 half-sib families. In each half-sib family, we mated one male (sire) to three females (dams). This design generates full siblings as well as half siblings. Although the initial design included 30 males and 90 females (3 females per male), only 67 females (distributed over 25 half-sib families) reproduced. In total, 1,228 eggs were produced. The offspring of each female (dam) was evenly split across a fully factorial combination of a manipulation of microbiome transmission (intact vs. disrupted) and larval brood ball modification (intact vs. disrupted).

**Figure S2:**
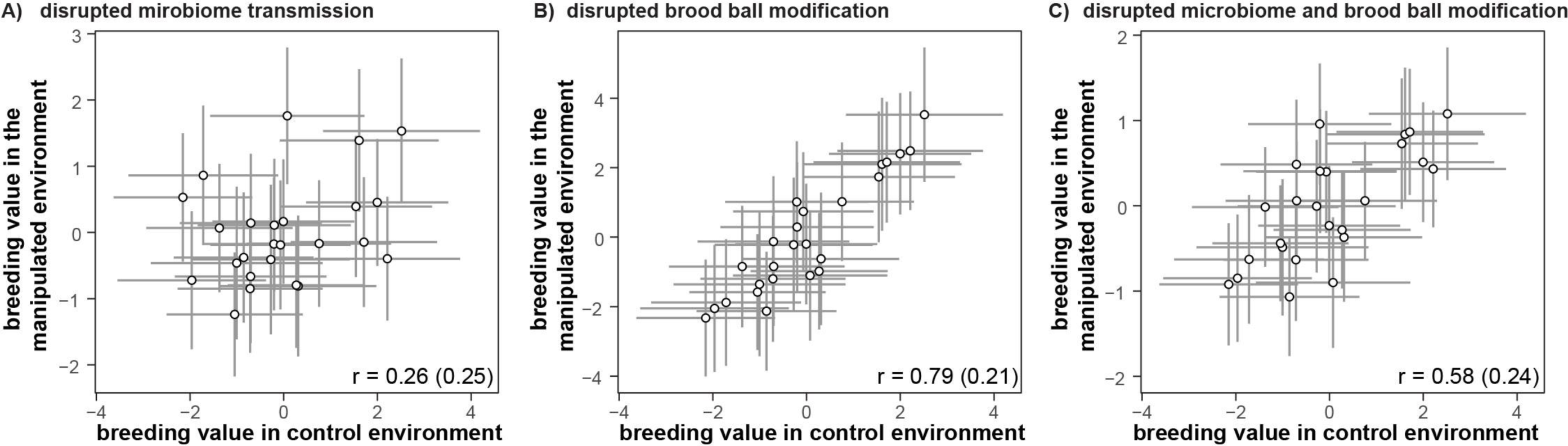
Correlation between a sire’s genetic value (plus SE) in the control treatment and the breeding values in treatments where microbiome nsmission and/or brood ball manipulations were disrupted (same values as shown in figure 4). Genetic correlations smaller than one indicate that es differ in the degree to which their genetic values are affected by the experimental manipulation. Breeding values (BLUPs), genetic correlations, corresponding standard errors were extracted from an animal model including all individuals (n = 932 offspring). Genetic correlations across atments were estimated using the ‘corgh’ variance structure (which allows to directly estimate and test genetic correlations in ASReml [48]). Residual variances were allowed to vary among treatments.

**Table S1:**
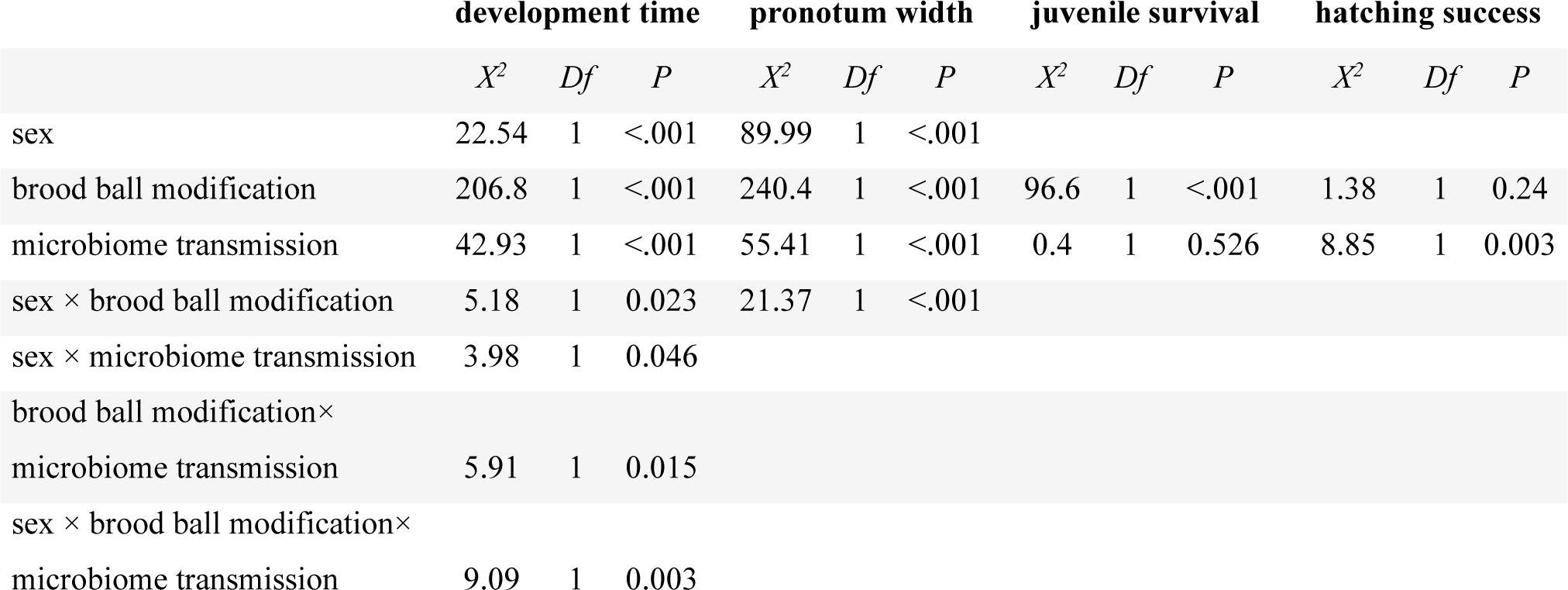
Analysis of Deviance Tables (Type II Wald Chi-square tests) for development time, pronotum width, juvenile survival, and hatching success. Juvenile survival and hatching success were fitted using generalized linear mixed models with a binomial error distribution. Because sex can only be determined in late larval development, sex could not be included in the models for juvenile survival and hatching success.

|  | development time |  |  | pronotum width |  |  | juvenile survival |  |  | hatching success |  |  |
| --- | --- | --- | --- | --- | --- | --- | --- | --- | --- | --- | --- | --- |
| | $X^2$ | $Df$ | $P$ | $X^2$ | $Df$ | $P$ | $X^2$ | $Df$ | $P$ | $X^2$ | $Df$ | $P$ |
| sex | 22.54 | 1 | <.001 | 89.99 | 1 | <.001 |  |  |  |  |  |  |
| brood ball modification | 206.8 | 1 | <.001 | 240.4 | 1 | <.001 | 96.6 | 1 | <.001 | 1.38 | 1 | 0.24 |
| microbiome transmission | 42.93 | 1 | <.001 | 55.41 | 1 | <.001 | 0.4 | 1 | 0.526 | 8.85 | 1 | 0.003 |
| sex × brood ball modification | 5.18 | 1 | 0.023 | 21.37 | 1 | <.001 |  |  |  |  |  |  |
| sex × microbiome transmission | 3.98 | 1 | 0.046 |  |  |  |  |  |  |  |  |  |
| brood ball modification ×<br>microbiome transmission | 5.91 | 1 | 0.015 |  |  |  |  |  |  |  |  |  |
| sex × brood ball modification ×<br>microbiome transmission | 9.09 | 1 | 0.003 |  |  |  |  |  |  |  |  |  |

